# ZetaHunter: a reproducible taxonomic classification tool for tracking the ecology of the Zetaproteobacteria and other poorly-resolved taxa

**DOI:** 10.1101/359620

**Authors:** Sean M. McAllister, Ryan M. Moore, Clara S Chan

## Abstract

Like many taxa, the Zetaproteobacteria lack well-defined taxonomic divisions, 32 making it difficult to compare between studies. We designed ZetaHunter to reproducibly 33 assign 16S rRNA gene sequences to previously-described OTUs based on a curated 34 database. While ZetaHunter can use any given database, we include a curated 35 classification of publically-available Zetaproteobacteria.

## Introduction

Taxonomic groups with limited cultivated representatives are often 38 lumped into a single taxonomic label encompassing multiple distinct ecological units. This makes it difficult to explicitly discuss the abundance, significance, and niche 40 preference of these units between studies. One group with poor taxonomic resolution is 41 the Zetaproteobacteria class, which is only accurately classified to this taxonomic level 42 by standard small subunit ribosomal RNA (16S rRNA) gene classification tools. ZetaHunter allows for reproducible, higher-resolution comparisons across studies by 44 using a curated dataset with a defined taxonomy based on operational taxonomic units 45 (OTUs). ZetaHunter comes with a curated database to identify the members of the Zetaproteobacteria, though it may be used with any curated 16S rRNA gene database.

## ZetaHunter implementation

ZetaHunter is a command line program written in Ruby, 49 designed to assign user-supplied 16S rRNA gene sequences to OTUs defined by a 50 reference sequence database. ZetaHunter can be used on Linux, MacOSX, and Windows 51 platforms through a Docker container, or through installation from source (Linux and MacOSX only). By default, ZetaHunter uses a curated database of Zetaproteobacteria 16S rRNA genes from ARB SILVA (release 128) and Zetaproteobacteria genomes from JGI’s Integrated Microbial Genomes (IMG) (see Supplemental Table 1). Zetaproteobacteria OTU (ZOTU) definitions include those reported by McAllister et al.(2011) at 97% identity, maintaining ZOTU number order from ZOTU1 to ZOTU28 for 57 ease in comparisons across studies. Numbered ZOTUs from ZOTU29 upward were 58 discovered after 2011. Sequences for use in ZetaHunter must be aligned using SINA (Pruesse et al., 2012).

The default pipeline of ZetaHunter (Figure 1) takes SINA-aligned 16S rRNA gene sequences and processes them as follows: 1) Input sequences are masked to the 1,282 bp used by McAllister et al. (2011). 2) Sequences are checked for chimeras using mothur’s uchime algorithm (Schloss et al., 2009; Edgar et al., 2011). 2) SortmeRNA is 64 used to cluster new sequences with the reference database, assigning ZOTU based on 65 genetic distance (closed reference binning; Kopylova et al., 2012). 3) The remaining 66 sequences are clustered into novel OTUs (NewZetaOtus) with mothur (*de novo* binning; 67 average neighbor; numbered by abundance) (Schloss et al., 2009). 4) Summary files (including final ZOTU calls and closest database hits), a biom table showing OTU 69 abundance per sample, and OTU network files are exported for use. The ZetaHunter 70 database includes 21 Proteobacteria outgroup sequences, allowing ZetaHunter to flag 71 sequences potentially outside the Zetaproteobacteria. Additionally, sequences that are 72 short, chimeric, or singletons are also flagged. This pipeline is an implementation of 73 open-reference OTU picking similar to the one found in QIIME (Navas-Molina et al., 2013).

**Figure 1.**
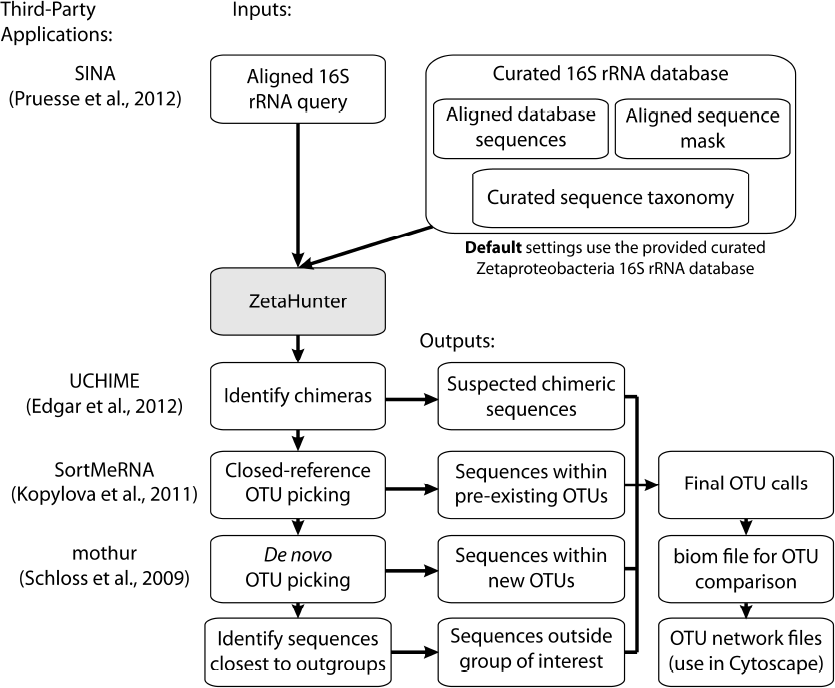
Flow chart showing the ZetaHunter pipeline. Third-party tools used in the 126 pipeline are indicated to the left.

To use ZetaHunter for non-Zetaproteobacteria, users need: 1) a database of SINA-aligned 16S rRNA gene sequences, 2) a sequence mask with asterisks at each informative 77 alignment column to be used for taxonomic assignment, and 3) a tab-delimited file 78 assigning each sequence to a particular taxonomic group at the similarity threshold 79 desired by the user.

## Software availability

ZetaHunter is available for download at <https://github.com/mooreryan/ZetaHunter>. GitHub documentation includes examples 83 for Zetaproteobacteria and non-Zetaproteobacteria classification and output file 84 explanations.

## Acknowledgements

We thank Dave Emerson for providing 16S rRNA sequences for the ZetaHunter database 88 from novel Zetaproteobacteria isolates. D Emerson and B Chiu are thanked for their 89 comments on early versions of this manuscript. We also thank K Hager, H Fullerton, J Scott, and J Vander Roost for testing and troubleshooting ZetaHunter using multiple 91 platforms and datasets. We also acknowledge the ARB/SILVA group for their work with 92 stable 16S rRNA alignments using the SINA tool; this tool is key to ZetaHunter’s 93 function. SMM and RMM designed the ZetaHunter program. SMM developed the Zetaproteobacteria database. RMM wrote the ZetaHunter program. SMM, RMM, and CSC wrote and edited this manuscript. This work was funded by NSF grant OCE-1155290 (to CSC) and USDA National Institute of Food and Agriculture award number 2012-68003-30155. Computational infrastructure support by the University of Delaware Center for Bioinformatics and Computational Biology Core Facility was made possible through funding from Delaware INBRE (NIH P20 GM103446) and the Delaware Biotechnology Institute. The authors declare no conflict of interest.

